# Epigenomic subtypes of late-onset Alzheimer’s disease reveal distinct microglial signatures

**DOI:** 10.1101/2025.03.15.643144

**Authors:** Valentin T. Laroche, Rachel Cavill, Morteza Kouhsar, Rick A. Reijnders, Joshua Harvey, Adam R. Smith, Jennifer Imm, Jarno Koetsier, Luke Weymouth, Lachlan MacBean, Giulia Pegoraro, Lars Eijssen, Byron Creese, Gunter Kenis, Betty M. Tijms, Daniel van den Hove, Katie Lunnon, Ehsan Pishva

## Abstract

Growing evidence suggests that clinical, pathological, and genetic heterogeneity in late-onset Alzheimer’s disease contributes to variable therapeutic outcomes, potentially explaining many trial failures. Advances in molecular subtyping through proteomic and transcriptomic profiling reveal distinct patient subgroups, highlighting disease complexity beyond amyloid-beta plaques and tau tangles. This insight underscores the need to expand molecular subtyping across new molecular layers, to identify novel drug targets for different patient subgroups.

In this study, we analyzed genome-wide DNA methylation data from three independent postmortem brain cohorts (n = 831) to identify epigenetic subtypes of late-onset Alzheimer’s disease. Unsupervised clustering approaches were employed to identify distinct DNA methylation patterns, with subsequent cross-cohort validation to ensure robustness and reproducibility. To explore the cell-type specificity of the identified epigenomic subtypes, we characterized their methylation signatures utilizing DNA methylation profiles derived from purified brain cells. Transcriptomic data from bulk and single-cell RNA sequencing were integrated to examine the functional impact of epigenetic subtypes on gene expression profiles. Finally, we performed statistical analyses to investigate associations between these DNA methylation-defined subtypes and clinical or neuropathological features, aiming to elucidate their biological significance and clinical implications.

We identified two distinct epigenomic subtypes of late-onset Alzheimer’s disease, each defined by reproducible DNA methylation patterns across three cohorts. Both subtypes exhibit cell-type-specific DNA methylation profiles. Subtype 1 and subtype 2 show significant microglial methylation enrichment, with odds ratios (OR) of 1.6 and 1.3, respectively. The minimal overlap between them suggests distinct microglial states. Additionally, subtype 2 displays strong neuronal (OR = 1.6) and oligodendrocyte (OR = 3.6) enrichment. Bulk transcriptomic analyses further highlighted divergent biological mechanisms underpinning these subtypes, with subtype 1 enriched for immune-related processes, and subtype 2 characterized predominantly by neuronal and synaptic functional pathways. Single-cell transcriptional profiling of microglia revealed subtype-specific inflammatory states: subtype 1 represented a state of chronic innate immune hyperactivation with impaired resolution, while subtype 2 exhibited a more dynamic inflammatory profile balancing pro-inflammatory signaling with reparative and regulatory mechanisms.

This study highlights the molecular heterogeneity of late-onset Alzheimer’s disease by identifying two epigenetic subtypes with distinct cell-type-specific DNA methylation patterns. Their alignment with previously defined molecular classifications underscores their relevance in disease pathogenesis. By linking these subtypes to inflammatory microglial activity, our findings provide a foundation for future precision medicine approaches in Alzheimer’s research and treatment.

## Introduction

Late-onset Alzheimer’s disease (LOAD) is the most common form of dementia, typically developing after the age of 65^1^. It primarily affects memory, cognitive function, and behavior. This condition is characterized by the gradual accumulation of amyloid-beta (Aβ) plaques and neurofibrillary tangles (NFTs) in the brain, resulting in the progressive destruction of neurons and brain atrophy^2^. However, significant heterogeneity has been observed among individuals with LOAD in terms of disease onset, progression, symptom variability, spreading of pathology, and their response to treatment^3–5^.

In recent years, molecular subtyping of AD has increasingly leveraged omics data to identify distinct molecular profiles that may contribute to the observed heterogeneity in disease manifestation and mechanisms. This has been particularly transformative as traditional classification methods based on clinical or pathological features alone have proven insufficient. A recent study that used mass spectrometry proteomics to analyze CSF protein profiles identified five distinct molecular subtypes of AD^6^. These subtypes highlight various underlying mechanisms, including neuronal plasticity, innate immune activation, RNA dysregulation, and dysfunction in the choroid plexus and blood-brain barrier. Another study, which examined the molecular heterogeneity of AD across multiple brain regions, identified five major AD subtypes using transcriptomic data^7^. Importantly, the immune-related and synaptic pathway subtypes predicted in the CSF proteomic are consistent with the transcriptomic-based subtypes of AD. Additionally, the transcriptomic findings revealed subtypes associated with protein metabolism and the upregulation of organic acid–related genes, further enriching our understanding of LOAD’s diverse molecular landscape.

Epigenetic modifications, such as DNA methylation (DNAm) and histone modifications, can alter gene expression without changing the DNA sequence. These modifications are influenced by both genetic predisposition and environmental factors, exerting diverse effects among individuals. Owing to these characteristics, epigenetic modifications serve as valuable molecular markers for identifying the drivers of disease heterogeneity. This is particularly relevant in the context of LOAD, where a complex interplay of genetic, lifestyle, and environmental factors are believed to contribute to the onset, progression, and variability of symptoms^1^. Moreover, an increasing body of evidence suggests that modifiable lifestyle factors, such as physical activity, may delay the onset of dementia^8^. Notably, our previous study highlighted that variation in methylation profile scores associated with lifestyle and environmental factors such as physical inactivity and low educational attainment as predictive markers for the prospective onset of dementia^9^. However, it remains unknown whether distinct DNAm subgroups exist within AD, which could provide further insights into disease heterogeneity.

In the present study, we used genome-wide DNAm data from three independent postmortem brain cohorts to identify epigenomic-based subtypes of LOAD. We employed data-driven methods to identify molecular subtypes of LOAD individuals based on the highest degree of similarity in methylomic patterns within each cohort. The subtypes were further validated through rigorous cross-cohort comparisons using multiple clustering algorithms. To better understand the complex biological mechanisms underlying the observed heterogeneity, we conducted a comprehensive characterization of predicted subtypes at cell-type-specific DNAm levels. Additionally, we examined AD risk genes as potential contributors to methylomic heterogeneity and identified transcriptomic correlates for each subtype using both bulk and single-cell RNA sequencing data.

## Materials and methods

### Brain samples

We analyzed 840 cortical postmortem brain samples obtained from three independent cohorts to investigate epigenomic-based heterogeneity in LOAD. We included 249 prefrontal cortex (PFC) samples from the UK Brain Banks Network (UKBBN), 220 PFC samples from the University of Pittsburgh’s Alzheimer’s Disease Research Center (PITT-ADRC)^10^, and 371 dorsolateral PFC (DLPFC) samples from the Religious Orders Study and Memory and Aging Project (ROSMAP)^11^. Samples were selected from donors with no or minimal AD pathology and definite AD (see Diagnostic Criteria). To further determine the brain cell-type specificity of the findings identified using bulk DNAm profiles, we isolated four nuclei populations from PFC tissue of 20 age and sex matched donors with low neuropathology from the Brains for Dementia Research (BDR) cohort^12^.

### Diagnostic Criteria

The diagnosis of LOAD and control samples was established based on clinical and neuropathological data from donors aged over 65 at the time of death. Dementia was defined by an antemortem Mini-Mental State Examination (MMSE) score below 24 or a Clinical Dementia Rating (CDR) score of ≥1. Within the dementia group, samples were classified as AD if they met the criteria of a "probable" or "definite" Consortium to Establish a Registry for Alzheimer’s Disease (CERAD) score for neuritic plaques and a Braak neurofibrillary tangle (NFT) stage of ≥3. In cases where CERAD data were unavailable, samples with a Braak stage of ≥5 were also included in the AD group.

Control samples were defined using stringent criteria specific to each brain bank. They were required to have a Braak stage of 0-2 for NFT pathology and negative for amyloid (CERAD neuritic plaque score 0). Additionally, when available from brain banks, both AD and control samples were screened for co-occurring other tau pathologies, including argyrophilic grain disease, as well as alpha-synuclein and TDP-43 pathologies. Only samples negative for these co-pathologies, with none to moderate cerebrovascular disease and rare microinfarcts, were included. Furthermore, samples from donors with known systemic hematopoietic malignancies, such as leukemia, were excluded to prevent the inclusion of malignant cells circulating in the brain.

### Methylomic profiling and data harmonization

For the UKBBN and PITT-ADRC cohorts, genomic DNA was extracted from 30 mg of fresh-frozen tissue using the AllPrep DNA/RNA/miRNA Universal Kit (QIAGEN), followed by bisulfite treatment with the EZ-96 DNA Methylation-Gold Kit (Zymo Research). The treated DNA was analyzed using Illumina’s Infinium MethylationEPIC v1.0 BeadChip arrays, quantifying DNAm at over 850,000 CpG sites. For the ROSMAP cohort, Illumina 450K methylation raw data files were obtained from the Accelerated Medicine Partnership (AMP-AD) portal (synID: syn7357283). Uniform and stringent quality control (QC) and normalization procedures were then applied across all three cohorts.

The raw methylation IDAT files were imported into R (version 4.3.0) using the ’readEPIC’ function from the wateRmelon R package (v.2.6.0)^13^, generating "MethylumiSet" objects. For quality control (QC), samples with a median signal intensity below 1000 were excluded. Additional outliers were removed using the ’outlyx’ function, which detects anomalies based on the Inter-Quartile Range (IQR) and Mahalanobis distance. Bisulfite conversion efficiency was assessed with the ’bscon’ function, discarding samples with efficiency below 0.8. Low-quality samples and probes were filtered using the ’pfilter’ function, removing samples with a detection P-value > 0.05 in more than 5% of probes and CpGs with fewer than three bead counts in 5% of samples or a detection P-value > 0.05 in 1% of samples.

Methylation sex was estimated using the ‘estimateSex’ function based on CpG sites on chromosomes X and Y, and any mismatches with true gender were excluded. Data normalization was performed using ’BMIQ’ (Beta-Mixture Quantile intra-sample normalization), preserving inter-sample variance. Probe-Wise Outlier Detection (PWOD) via the ’homonymous’ function removed probes associated with known variants to prevent downstream biases. Additional filtering excluded probes with SNP IDs, cross-hybridizing probes^14^, and those on sex chromosomes.

Cell type composition in bulk PFC DNAm data was computed using CETYGO (v0.99.0)^15^, employing a reference panel to distinguish inhibitory (GABAergic) neurons, excitatory (glutamatergic) neurons, oligodendrocytes, microglia, and astrocytes. To account for mutual technical and biological effects (e.g., age, sex, postmortem interval, plates, cell composition, and cohort-specific batches), a linear multiple regression model was applied to each methylation probe. Non-variant probes were filtered using the 90th percentile absolute deviation, resulting in 325,834 CpGs from EPIC arrays for subtyping. In the ROSMAP cohort, the CpG set was defined by the overlap between the processed 450K array and the available 450K probes within the final EPIC CpGs, yielding 148,951 CpGs.

Following pre-processing and QC, the final dataset included 240 UKBBN samples (129 LOAD, 111 controls), 220 PITT-ADRC samples (192 LOAD, 28 controls), and 371 ROSMAP samples (248 LOAD, 123 controls). This dataset, detailed in **Supplementary Table 1**, integrates diverse demographics, clinical profiles, and neuropathological information.

### Cell type-specific DNA methylation profiling

For the BDR samples, nuclei isolation was performed using the Fluorescence-Activated Nuclei Sorting (FANS) method^16^. A FACS Aria III cell sorter (BD Biosciences) was used to simultaneously collect cell populations labeled with immunostaining markers: NeuN, SOX10, and IRF8. The following four nuclei populations were isolated: neurons (NeuN ), oligodendrocytes (NeuN /SOX10 ), microglia (NeuN /SOX10 /IRF8 ), and astrocytes (NeuN /SOX10 /IRF8 ). A detailed protocol for nuclei purification is available at https://www.protocols.io/view/fluorescence-activated-nuclei-sorting-fans-on-huma-36wgq4965vk5/v1. DNA isolation and bisulfite conversion were performed using the EZ DNA Methylation Direct™ Kit (Zymo Research) for each nuclei population, followed by DNAm quantification using Illumina EPIC (v1) arrays.

### Bulk RNA sequencing

Total RNA quality in the UKBBN and PITT-ADRC cohorts was assessed using the Agilent 4200 TapeStation System, with samples having an RNA Integrity Number (RIN) below 3 excluded. Library preparation was conducted using the Illumina Stranded mRNA Prep kit, followed by sequencing on the NovaSeq 6000 platform at the Exeter Sequencing Service.

Raw sequencing data in FASTQ format underwent quality trimming to remove low-quality bases and adapter sequences. Processed reads were aligned to the human reference genome (GRCh37) using the STAR aligner (v2.7.3a) with default parameters^17^. After alignment, a raw count matrix was generated, and genes with low expression, defined as counts below 10 in over 80% of samples. Gene counts were normalized using the Trimmed Mean of M-values (TMM) method from the edgeR package (v3.42.4)^18^. The normalized counts were then transformed into log-transformed counts per million (logCPM) to stabilize variance and improve interpretability. Following quality control and processing, the RNA sequencing dataset comprised 483 cortical samples from three independent postmortem brain cohorts: UKBBN, PITT-ADRC, and ROSMAP. The dataset included 174 samples classified as LOAD-S1, with 63 from UKBBN, 57 from PITT-ADRC, and 54 from ROSMAP. LOAD-S2 consisted of 107 samples, with 45 from UKBBN, 20 from PITT-ADRC, and 42 from ROSMAP. The control group contained 202 samples, including 109 from UKBBN, 25 from PITT-ADRC, and 68 from ROSMAP.

### Genotyping, imputation, and generation of polygenic scores

Genotyping was conducted using the Illumina Global Screening Array (GSA) for the UKBBN and PITT-ADRC cohorts. For the ROSMAP cohort, genotyping data from the Affymetrix GeneChip 6.0 (Affymetrix, Inc.) and the Illumina HumanOmniExpress chip array were obtained from (https://www.synapse.org/ ). Quality control was performed using PLINK (v2.0), excluding samples with >5% missing values and SNPs with >1% missing values. Additionally, SNPs with a minor allele frequency (MAF) < 0.05 and a Hardy-Weinberg equilibrium p-value < 1.0 × 10 were filtered out to retain the most reliable signals.

Processed genotype data were merged with the 1000 Genomes Project dataset, and principal component analysis (PCA) was performed to determine sample ethnicity. Samples of non-European ancestry were removed based on PCA results. The quality-controlled genotype data were uploaded to the Michigan Imputation Server and imputed using Minimac4 with the 1000 Genomes reference panel (phase 3, version 5). The imputed data for each chromosome were compiled into a unified VCF file using BCFtools (v1.9)^19^ with tri-allelic SNPs removed using VCFtools (v0.1.16)^20^. The final VCF file was converted into PLINK binary format.

Polygenic scores (PGS) were generated using the imputed genotype data and PRSice-2 software.^21^. Genetic variants associated with AD were selected based on the genome-wide association study (GWAS) by Bellenguez et al., 2022 ^22^.

## Data analysis

### Clustering algorithms

To identify cortical brain subgroups with similar DNAm profiles, agglomerative hierarchical clustering using the Ward D2 method with Euclidean distance and K-means clustering as a non-hierarchical approach were applied to LOAD samples, with each cohort analyzed independently. The optimal number of clusters for both methods was determined using the Elbow method.

To assess clustering consistency, Normalized Mutual Information (NMI) analysis was performed for each cohort using the Aricode R package (v1.0.3)^23^ (https://github.com/jchiquet/aricode), comparing hierarchical and K-means clustering results. The cluster number with the highest NMI value was selected for further analysis.

To assess cluster specificity, Sparse Partial Least Squares Discriminant Analysis (sPLS-DA) was performed using the mixOmics package (v6.24.0)^24^ for each clustering algorithm to identify the most discriminative features. From each sPLS-DA model, 12,000 features corresponding to six components were extracted. To evaluate robustness, cluster labels were randomized 10 times across population proportions ranging from 100% to 1%, followed by repeated sPLS-DA modeling for each iteration. The overlap between the 12,000 extracted features from each iteration and those from the original cluster labels was then examined.

Technical batch effects, including plate and chip variations, were assessed using Spearman’s correlation test for continuous variables and Normalized Mutual Information (NMI) for discrete variables, the latter being particularly suited for evaluating clustering outcomes.

To verify the generalizability of findings across Illumina EPIC arrays (UKBBN and PITT-ADRC cohorts) and 450K arrays (ROSMAP cohort), Weighted Gene Co-expression Network Analysis (WGCNA) was employed. Co-expression networks were constructed using the ’blockwiseModules’ function in the WGCNA R package (v1.72-5)^25^ based on selected EPIC array probes. Module eigenprobes were then calculated using the ’moduleEigengenes’ function, and their associations with categorical cluster labels were assessed through ANOVA followed by post hoc Tukey’s tests. For modules showing significant differences between identified clusters, eigenprobes were recalculated using the subset of probes available on the 450K array. Cluster-associated co-methylated probe preservation was confirmed by demonstrating that eigenvalue differences between cluster labels remained significant with the 450K probes and that eigenvalue differences for specific subtypes across both platforms were non-significant.

### Cross-cohort replication

To identify matching clusters with maximum similarity in DNAm profiles across the UKBBN, PITT-ADRC, and ROSMAP cohorts, we employed two complementary approaches at both the probe and sample levels.

In the first approach, we calculated the median DNAm value per probe for each clustered brain sample within each cohort. Pearson correlation tests were then used to assess the relationship between these median values across the three cohorts.

In the second approach, the UKBBN cohort was designated as the discovery dataset. Using the ‘splsda’ function from the mixOmics package (v6.24.0), we reduced data into a lower-dimensional latent space while preserving class separation. The first two latent spaces of sPLS-DA were mapped against the clusters identified in this cohort. Convex hulls were established around each UKBBN cluster by connecting the outermost points. DNAm profiles from a replication cohort (either PITT-ADRC or ROSMAP) were then projected onto these latent spaces, and the number of replication cohort samples falling within the UKBBN cluster hulls was recorded. To assess significance, contingency tables were constructed, and a hypergeometric test was performed to determine overlaps between the discovery and replication cohort clusters. This procedure was repeated with PITT-ADRC and ROSMAP serving as discovery sets, while the other two cohorts acted as replication cohorts. An epigenomic-based LOAD subtype was confirmed only when both approaches validated the DNAm profile similarities across all three cohorts, at both the probe and sample levels, using two clustering methods.

### Subtype-specific epigenome-wide association analysis

Epigenome-wide association studies (EWASs) were conducted using multiple linear regression models to identify differentially methylated positions (DMPs) associated with each predicted subtype in each cohort. Methylation data were rigorously adjusted for potential confounders, including age, sex, cell type composition, brain banks, and plates, ensuring consistency with previous clustering steps. Surrogate variable analysis (SVA) was applied to adjust for unwanted variations unrelated to the main outcome, further refining accuracy. P-value inflation was assessed using the inflation index for each EWAS and estimates and standard errors were corrected using the Bacon package in R (v1.28.0)^26^ before proceeding to the meta-analysis. The meta-analysis was performed using the inverse variance method via the rma.uni function in the metafor R package (v4.6-0)^27^. Subtype-specific DNAm signatures were identified using relaxed criteria. Specifically, DMPs were required to meet the following criteria: a p-value < 1.0 × 10 ³, the same direction of estimates across all three studies, and an absolute Cohen’s D > 0.2 in at least one cohort. The overlap between DMPs identified across subtypes in the meta-analysis was assessed using the geneOverlap R package (v1.40.0) (http://shenlab-sinai.github.io/shenlab-sinai/).

### Cell-type specific DNAm enrichment analysis

Cell-type-specific EWASs were conducted using generalized mixed models, comparing each cell type against all others to establish distinct DNAm profiles. Significance criteria were set at a nominal p-value < 1.0 × 10 ¹ and absolute effect estimates > 0.5. The overlap between differentially methylated positions (DMPs) identified in the meta-analyses across subtypes and cell-type-specific DNAm signatures was assessed using the geneOverlap R package (v1.40.0).

### Colocalization analysis

Methylation quantitative trait loci (mQTL) data from ROSMAP were obtained from Brain xQTLServe (https://mostafavilab.stat.ubc.ca/xqtl/). Cis-mQTLs (p ≤ 1.0 × 10 ) associated with differentially DMPs identified in the subtype-specific EWAS were extracted. Bayesian colocalization analysis was performed using the coloc R package (v5.2.3)^28^ for all pairwise combinations of genomic regions linked to AD, as identified by Bellenguez et al.^22^ and variants associated with LOAD subtype DMPs. The ‘coloc.abf’ function was used for the analysis, with colocalizations confirmed when the combined probability of H3 and H4 exceeded 0.95.

### Bulk transcriptomic analysis

Differential expression analysis (DEA) was performed using the Limma R package (v3.55.10)^29^, comparing LOAD subtypes to their respective controls across the three cohorts. To account for potential confounders, the analysis included covariates such as age, sex, RIN, brain banks, and surrogate variables, which were incorporated to adjust for unmeasured variation, including differences in cell type composition. Bacon correction was applied to estimates and standard errors to reduce bias. Genes were considered significantly differentially expressed if they showed a nominal p-value below 0.05 in all cohorts, a fold change greater than 1.5 in at least one cohort, and a consistent direction of effect across all cohorts.

### Single-cell transcriptomic analysis

Analysis of gene expression across 12 microglial states was conducted using single-cell RNA sequencing data from the ROSMAP study. This dataset comprises 427 samples, including 70 definite AD cases and 64 control samples, overlapping with the subset of ROSMAP data utilized in this project. Subtype-specific differentially expressed genes (DEGs) were identified for each of the 12 microglial states characterized by Sun et al.^30^

The muscat R package (v1.14.0) was used with default parameters. Non-expressed or lowly expressed genes were filtered out, and outlier detection was performed at the cell level. To generate a total expression value for each individual, cells were aggregated in a pseudobulk manner by summing RNA expression from all cells within each microglial state. Pseudobulk differential DEA was then conducted for each microglial state, comparing subtype samples to healthy controls.Genes were considered significantly differentially expressed if they met the criteria of a nominal p-value < 0.01 and fold change (FC) > 1.5.

### GO enrichment analysis

GO enrichment analysis was conducted using the clusterProfiler R package (v4.12.6) ^31^ to identify biological processes, molecular functions, and cellular components significantly associated with differentially expressed genes (DEGs) for each subtype. For subtype-specific microglial transcriptomic signatures, enrichment analysis focused exclusively on GO terms related to microglial activity, immune responses, and pro- and anti-inflammatory states^32^.

## Results

### Data-driven clustering

The clustering analysis was conducted on 325,834 CpGs from EPIC arrays and 148,951 CpGs from 450K arrays, following stringent preprocessing and filtering steps (see Methods). Clustering algorithms were applied exclusively to samples with a definite LOAD diagnosis, initially identifying three to five optimal clusters across the three cohorts when evaluated using the Elbow method **(Supplementary Fig. 1**).

Further validation with the NMI test demonstrated high similarity between hierarchical and K-means clustering methods, with the strongest agreement observed when selecting three clusters (**Supplementary Table 2**). To maintain consistency across cohorts, clusters within each dataset were randomly labeled as A, B, and C.

The specificity of the detected clusters was further validated by randomizing cluster labels across varying proportions of the sample populations. Minimal overlap was observed between the features distinguishing the original clusters and those identified under randomized labels, particularly when randomization was applied to the entire dataset. This finding indicates that the original clusters are distinct and unlikely to result from random assignment, reinforcing the robustness and biological relevance of the detected clusters (**Supplementary Fig. 2**).

No significant association was observed between the identified clusters and known technical batch effects, including plate variations and brain sample sources (**Supplementary Table 3**). This confirms the reliability of the clustering results, suggesting that the detected clusters more likely reflect true biological differences rather than procedural inconsistencies. The distribution of samples assigned to each cluster using both clustering methods across the three cohorts is reported in **Supplementary Table 4**.

Illumina EPIC arrays were used for DNAm quantification in the UKBBN and PITT-ADRC cohorts, covering over 850K CpG sites, while the ROSMAP cohort utilized the 450K array. To assess the generalizability of findings across these two array types, weighted gene co-expression network analysis (WGCNA) was applied. The significant relationship between module eigenvalues and the three identified clusters, as determined by ANOVA, remained preserved when reconstructing modules using only the subset of CpGs available on 450K arrays. Additionally, no significant changes in module eigenvalues within the same cluster were observed when transitioning from EPIC to 450K arrays, supporting the robustness and cross-platform reproducibility of the clustering results (**Supplementary Figs. 3–6**).

### Cross-cohort replication of epigenomic-based subtypes

Cross-cohort replication analyses identified two distinct subtypes of LOAD based on DNAm profiles measured in bulk cortical brain samples. Replication was conducted using two complementary strategies at the CpG and sample levels.

CpG-level replication assessed the correlation between median DNAm values per CpG for each data-driven cluster across the three cohorts (see Methods). This analysis revealed two distinct blocks of correlated clusters: the first block included UKBBN cluster A, PITT-ADRC cluster B, and ROSMAP cluster A, while the second block comprised UKBBN cluster B, PITT-ADRC cluster C, and ROSMAP cluster B. A detailed heatmap plot illustrating the associations between clusters using both methods is provided in **Fig. 1a**.

**Figure 1.**
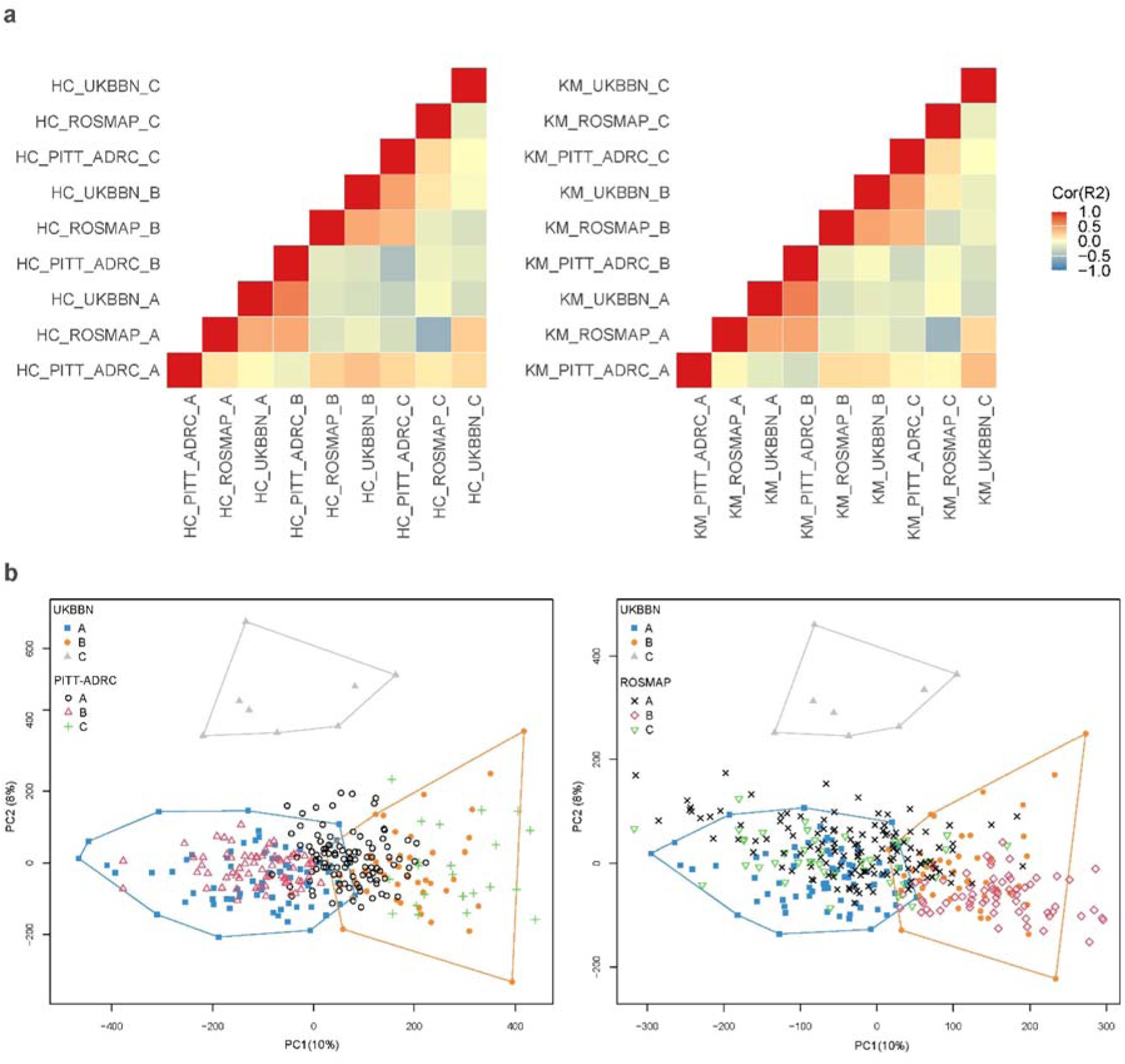
Cross-cohort generalizability of DNAm-based clusters. **(a)** Visual representation of the correlation-based replication of the identified clusters across three cohorts using Hierarchical (Left), and K-means (Right) methods. The first block (LOAD-S1) includes UKBBN cluster A, PITT-ADRC cluster B, and ROSMAP cluster A, while the second block (LOAD-S2) includes UKBBN cluster B, PITT-ADRC cluster C, and ROSMAP cluster B. **(b)** Spatial overlap analysis showing cluster assignments across the three cohorts. In this example, the clusters were confirmed through iterative projections, where UKBBN cohort’s first two latent spaces were projected onto the PTT-ADRC and ROSMAP to verify spatial overlap across datasets.

The second replication analysis examined the spatial overlap of the distinct clusters across the three cohorts using latent spaces. Matching clusters across independent cohorts were confirmed by iteratively swapping the study samples, where the latent spaces from one cohort were used to project the other two (see Methods). Notably, the two blocks of clusters identified through correlation analysis were further validated, showing significant overlap in latent spaces, as assessed by Fisher’s tests. An example projection of UKBBN latent spaces onto PITT-ADRC and ROSMAP datasets is illustrated in **Fig. 1b**, with detailed Fisher’s test statistics provided in **Supplementary Table 5**.

In both analyses, no matching clusters were identified for UKBBN cluster C, PITT-ADRC cluster A, and ROSMAP cluster C. As a result, these clusters were labeled as ’Unassigned’ samples for further analysis. This classification remained consistent across both hierarchical and K-means clustering methods, reinforcing the robustness of the clustering results across different methodologies. To ensure subtype consistency, only samples consistently labeled as a subtype across both clustering methods were retained, with the final counts for each subtype reported in **Supplementary Table 6**. From this point forward, the first block of clusters is referred to as Subtype 1 (LOAD-S1) and the second block as Subtype 2 (LOAD-S2). **Fig. 2a** presents a plot of the first two principal components (PCs) for the pooled dataset comprising UKBBN, PITT-ADRC, and ROSMAP cohorts, with samples labeled according to their confirmed LOAD subtypes.

**Figure 2.**
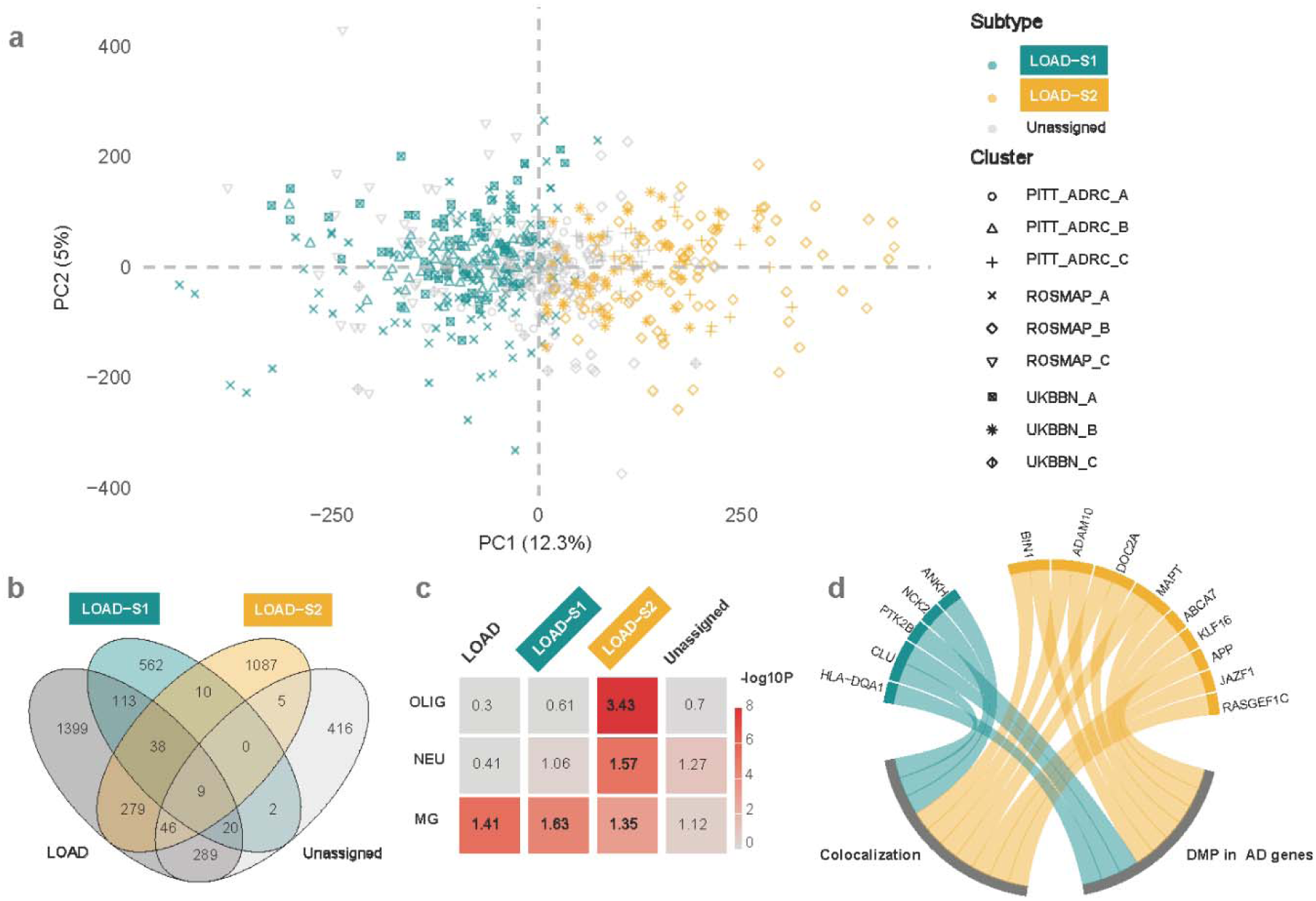
Characterization of the methylomic signatures of predicted LOAD subtypes. **(a)** Visualization of first two PCs for LOAD subtypes across all cohorts. Plot of the first two PCs (PC1 and PC2) for DNAm profiles from all samples in the UKBBN, PITT-ADRC, and ROSMAP cohorts. Samples are labeled according to their subtype assignments: Subtype 1 (LOAD-S1) and Subtype 2 (LOAD-S2). ’Unassigned’ samples are also indicated, representing clusters without matching correlations across cohorts. **(b)** Venn diagram illustrating the overlap of differentially methylated positions (DMPs) associated with LOAD-S1, LOAD-S2, and the Unassigned group relative to overall LOAD. Minimal overlap is observed between LOAD-S1 and LOAD-S2, highlighting their distinct methylomic profiles. **(c)** CpG overlap of the cell type-specific methylation signatures for overall LOAD, LOAD-S1, LOAD-S2 and Unassigned group derived from FANS data. LOAD-S1 shows a stronger association with microglial DMPs, whereas LOAD-S2 displays significant contributions from neuronal, oligodendrocyte, and microglial DNAm profiles. **(d)** Subtype-specific methylation quantitative trait loci (mQTL) mapping and genomic analysis. Key subtype-specific CpG sites annotated to AD-related genes are shown (Right).

### Subtype-specific EWAS

The distinct methylomic signatures of the newly defined LOAD subtypes (LOAD-S1 and LOAD-S2), the Unassigned group, and overall LOAD were characterized using EWAS) across the UKBBN, PITT-ADRC, and ROSMAP cohorts, followed by meta-analyses. These analyses focused on 148,951 highly variable CpGs shared between the EPIC and 450K arrays, which were also used in the clustering algorithms (see Methods). Comparing DNAm profiles of healthy controls (HC) with overall LOAD identified 2,193 DMPs. Additionally, 697 DMPs were associated with LOAD-S1, 1,147 with LOAD-S2, and 787 with the Unassigned group. DMPs were considered significant if they met the criteria of p-value < 1.0 × 10 ³ and exhibited a medium to large effect size (absolute Cohen’s D > 0.2) in at least one cohort, with a consistent direction of effect across all three cohorts (**Supplementary Tables 7–10**).

LOAD-S1 and LOAD-S2 exhibited distinct DNAm profiles, with minimal overlap in their DMPs (odds ratio [OR] = 8.46). The greatest overlap with overall LOAD DMPs was observed in the Unassigned group (OR = 68.82), followed by LOAD-S2 (OR = 26.98) and LOAD-S1 (OR = 22.76). These findings highlight that LOAD-S1 and LOAD-S2 represent the most distinct DNAm profiles, particularly in comparison to the Unassigned group. Similarities and differences among LOAD subtypes and the Unassigned group relative to overall LOAD DMPs are detailed in **Supplementary Table 11**, with CpG overlaps visualized in **Fig. 2b**.

To evaluate the relationship between subtype-specific methylomic signatures identified in this study and previously reported DMPs associated with AD pathology, we examined the overlap of significant DMPs linked to neurofibrillary tangles (NFTs) in the PFC as reported by our group^33^. Of the 236 Bonferroni-significant DMPs linked to AD pathology in that study, 150 CpGs were included in the subset analyzed in the current study. The greatest overlap was observed with overall LOAD (128 CpGs; OR = 84.25) and the Unassigned group (41 CpGs; OR = 41.69). Notably, LOAD-S2 displayed a greater overlap with previously reported DMPs from the PFC DNAm meta-analysis (48 CpGs; OR = 26.36) compared to LOAD-S1 (14 CpGs; OR = 12.61). These findings suggest that LOAD-S2 is more strongly associated with AD pathology than LOAD-S1 (**Supplementary Table 12**).

### Cell type-specific methylation and LOAD subtypes

To investigate the relationship between LOAD subtypes and brain cell types, we characterized differentially methylated signatures across four cell types—neurons, oligodendrocytes, microglia, and astrocytes using DNAm data from purified brain cerlls of 20 healthy postmortem PFC samples (**Supplementary Table 13**; see Methods). Overlap analyses of LOAD, LOAD-S1, LOAD-S2, and Unassigned group-associated DMPs within these cell-type-specific methylation signatures revealed a significant enrichment of LOAD-S1 and LOAD-S2 DMPs in microglia, with ORs of 1.6 and 1.3, respectively. Notably, the overlap between the microglial methylation signatures of LOAD-S1 and LOAD-S2 was minimal, with only 1.2% of LOAD-S1 DMPs and 0.6% of LOAD-S2 DMPs overlapping, suggesting distinct microglial signatures despite their shared association with this cell type.

In addition to microglial involvement, LOAD-S2 DMPs were also significantly enriched in neuronal (OR = 1.6) and oligodendrocyte (OR = 3.6) DNAm signatures (**Fig. 2c; Supplementary Table 14**). Conversely, DMPs associated with the Unassigned group showed no enrichment in the DNAm signatures of any examined cell types, underscoring the specificity of the LOAD-S1 and LOAD-S2 methylation signatures.

### AD genetics and epigenomic-based subtypes

To investigate the contribution of AD genetic risk variants to the epigenomic-based heterogeneity in LOAD, we examined AD risk genomic regions in relation to subtype-specific DMPs. By mapping these subtype-specific DMPs to mQTL data from the ROSMAP database, we identified 18,577 cis-mQTLs associated with LOAD-S1 and 31,045 cis-mQTLs associated with LOAD-S2.

Bayesian colocalization analysis revealed that methylation loci in these subtypes are influenced by distinct AD-associated genomic regions across different chromosomes. Specifically, LOAD-S1 showed colocalization with genetic regions linked to *HLA-DQA1*, *CLU*, and *PTK2B*, while LOAD-S2 methylation signatures were associated with variants in *ADAM10*, *DOC2A*, *MAPT*, *ABCA7, BIN1*, and *KLF16* (**Fig. 2d**).

Further analysis of CpGs linked to distinct AD risk genes revealed significant methylation changes at one or more CpG sites within specific risk genes for each subtype. CpGs annotated to *CLU*, *ANKH*, and *NCK2* were specific to LOAD-S1, while those annotated to *SPL1*, *RASGEF1C*, *MAPT, JAZF1, DOC2A, BIN1, APP*, and *ADAM10* were among the significant DMPs specific to LOAD-S2 (**Fig. 2d**).

### Distinct transcriptomic profiles and pathway enrichment in LOAD subtypes

Analysis of bulk PFC RNA-sequencing data identified 162 genes in LOAD-S1 and 277 in LOAD-S2 (nominal p-value < 0.05 in all three cohorts, FC > 1.5 in at least one, with a consistent direction of effect). (**Supplementary Tables 14-15**).

GO enrichment analysis revealed that LOAD-S1 is driven by immune dysregulation and inflammation, with enriched terms including ‘cell activation in immune response’ (GO:0002263), ‘positive regulation of cytokine production’ (GO:0001819), and ‘tumor necrosis factor production’ (GO:0032640). In contrast, LOAD-S2 was enriched in synaptic and neuronal processes, including ‘vesicle-mediated transport in synapse’ (GO:0099003), ‘inhibitory synapse assembly’ (GO:1904862), and ‘neurotransmitter transport’ (GO:0006836). These findings highlight distinct molecular pathways driving LOAD subtypes, reinforcing its biological heterogeneity (**Supplementary Tables 16–17, Fig. 3a**).

**Figure 3.**
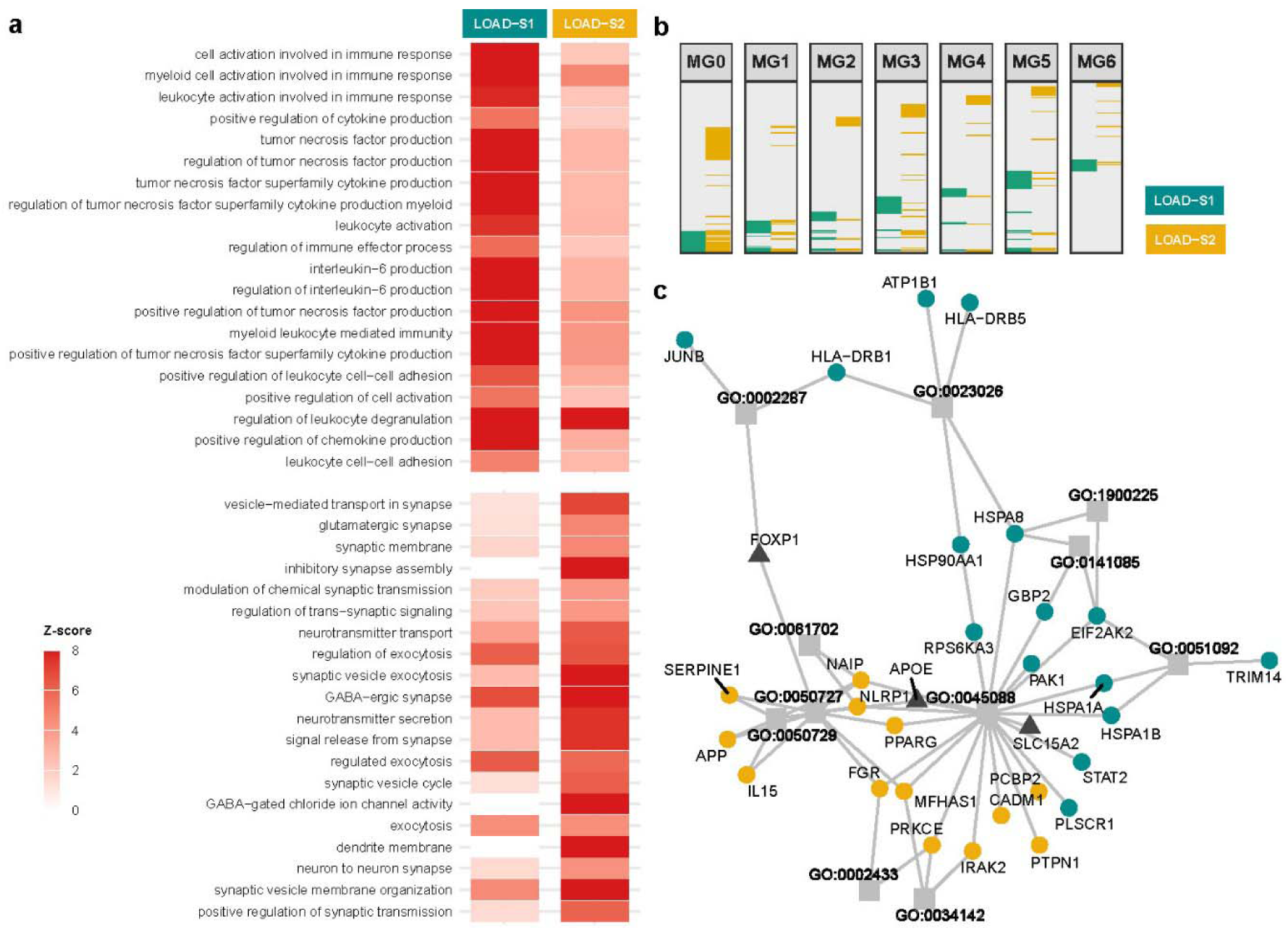
Transcriptomic characterization of LOAD epigenomic Subtypes. **(a)** Heatmap showing Z-scores for the top 20 enriched Gene Ontology (GO) terms in LOAD-S1 and LOAD-S2 subtypes. Z-scores indicate the deviation of observed gene counts from expected values, normalized by standard deviation. LOAD-S1 is enriched in immune-related pathways, reflecting a dominant immune/inflammatory signature, while LOAD-S2 shows enrichment in synaptic and neuronal pathways. Immune pathways prevalent in LOAD-S1 are demonstrating weak enrichment in LOAD-S2, and vice versa. **(b)** Comparative analysis of differentially expressed genes (DEGs) across seven microglial (MG) states (MG 0–6) highlights distinct gene expression profiles for LOAD-S1 and LOAD-S2, with minimal overlap, underscoring subtype-specific transcriptional landscapes. **(c)** GO enrichment analysis of microglial single-cell DEGs reveals shared enrichment in the ’regulation of innate immune response’ (GO:0045088) pathway, but with distinct contributing genes. LOAD-S1-specific pathways include NF-kappaB regulation (GO:0051092), inflammasome signaling (GO:0141085), NLRP3 assembly (GO:1900225), MHC class II binding (GO:0023026), and T cell activation (GO:0002287). LOAD-S2-specific pathways include TLR4 signaling (GO:0034142), receptor signaling in phagocytosis (GO:0002433), and inflammatory response regulation (GO:0050727, GO:0050729, GO:0061702).

### Subtype-specific microglial transcriptomes and immune profiles

Analysis of single-cell RNA sequencing data from the ROSMAP study explored the distinct microglial signatures associated with LOAD subtypes at the methylation level. This analysis focused on 12 microglial (MG) states, as characterized by Sun et al.^30^ and was conducted by aggregating single-cell data into pseudobulk profiles for each MG state and subtype. DEA was performed on seven MG states (MG 0–6) that had sufficient cell numbers and read depths for both subtype and control samples. Across these seven states, 199 DEGs were identified in LOAD-S1 and 220 in LOAD-S2 (nominal p-value < 0.01, FC > 1.5) (**Supplementary Tables 19–20**). Comparative analysis revealed minimal overlap in DEGs across MG states, with an average overlap of 10.82% (SD = 5.18) (**Fig. 3b**).

GO enrichment analysis focused on microglial activity, immune responses, and pro- and anti-inflammatory states listed in **Supplementary Table 21**. The results revealed both shared and distinct immune-related processes across LOAD subtypes. Genes from both subtypes were enriched in the ‘regulation of innate immune response’ pathway (GO:0045088). However, the specific genes contributing to this pathway varied significantly between LOAD-S1 and LOAD-S2. In LOAD-S1, enriched genes included *HSPA8, HSPA1A, HSPA1B, HSP90AA1, APOE, GBP2, SLC15A2, PLSCR1, EIF2AK2, STAT2, RPS6KA3*, and *PAK1*. In contrast, LOAD-S2 was characterized by genes such as *FGR, PRKCE, APOE, MFHAS1, SLC15A2, CADM1, PPARG, NLRP1, PCBP2, PTPN1, IRAK2*, and *NAIP*. These findings highlight commonalities and subtype-specific variations in immune-related mechanisms in LOAD (**Fig. 3c**). A detailed list of significant GO terms and the genes contributing to each pathway can be found in **Supplementary Tables 22–23.**

### Clinical and demographic characterization of LOAD subtypes

Pathological and clinical assessments revealed no significant differences between LOAD-S1 and LOAD-S2 in terms of APOE status, polygenic risk scores for AD, age of onset, last cognitive assessment, or key measures of NFT and amyloid pathology (**Supplementary Table 24**). These results suggest that the distinct epigenetic and transcriptomic signatures defining the subtypes, particularly their enrichment in specific cell-type DNAm patterns, operate independently of traditional clinical and pathological markers of AD.

## Discussion

This study identified two distinct epigenetic subtypes of LOAD by analyzing genome-wide DNAm profiles across three large-scale, independent postmortem brain cohorts. While these subtypes showed no association with established clinicopathological markers of LOAD, cellular and molecular characterization revealed distinct DNAm and transcriptional profiles, highlighting their biological relevance and potential role in LOAD heterogeneity.

LOAD-S1 exhibited enriched DNA methylation signatures in microglia, indicating a predominantly immune and inflammatory association. In contrast, LOAD-S2 displayed methylation enrichment in microglia, neurons, and oligodendrocytes, suggesting a broader cellular involvement. Consistent with these patterns, bulk RNA sequencing analysis revealed that LOAD-S1 is primarily linked to immune and inflammatory pathways, whereas LOAD-S2 showed stronger associations with synaptic dysfunction and neuronal communication, highlighting distinct molecular mechanisms underlying these subtypes.

Single-cell RNA sequencing analysis of microglial states further revealed unique immune-related gene expression patterns driving LOAD-S1 and LOAD-S2. While both subtypes exhibit enrichment in innate immune response pathways, their underlying gene sets diverge significantly, suggesting distinct immune activation, regulation, and resolution mechanisms. LOAD-S1 is characterized by heightened innate immune activation and impaired inflammatory resolution, potentially contributing to neurotoxicity and disease progression. Upregulated *STAT2* and *RPS6KA3* promote NF-κB signaling and cytokine production, sustaining a pro-inflammatory state^34,35^. The increased expression of *EIF2AK2* indicates heightened antiviral and interferon-stimulated gene expression, which sustains immune signaling pathways^36^. Additionally, increased expression of *GBP2* suggests heightened inflammasome activity and IL-1β secretion, further amplifying inflammatory signaling^37^. Meanwhile, downregulation of *HSPA8*, *HSPA1A/B*, and *HSP90AA1* suggests impaired cellular stress responses and protein stabilization, reducing the ability to resolve inflammation effectively.^38^. Collectively, these changes in LOAD-S1 suggest increased innate immune activation, impaired inflammatory resolution, and a sustained pro-inflammatory state, which may drive neurotoxicity and accelerated disease progression.

LOAD-S2, in contrast, presents a more dynamic immune profile, balancing pro-inflammatory and reparative signals. Upregulated *FGR* and *APP* enhance innate immune receptor signaling and cytokine production, sustaining immune activation^39,40^. Increased IL15 expression supports NK and T cell activation and survival, further sustaining immune responses^41^. However, downregulation of *NLRP1* suggests reduced inflammasome activation, potentially limiting IL-1β-mediated neurotoxicity^42^. While *APP* upregulation maintains inflammatory signaling, *NAIP* upregulation may counter-regulate excessive immune activation, preventing immune damage^43^. Additionally, upregulated *SERPINE1* suggests active immune regulation via lipid metabolism and fibrinolysis, contributing to a more balanced immune response in LOAD-S2 compared to LOAD-S1^44^.

Additionally, distinct genomic regions associated with AD may contribute to the methylomic heterogeneity observed in LOAD subtypes. AD genes linked to LOAD-S1 are primarily associated with immune regulation (*HLA-DQA1*)^45^, amyloid-beta processing (*CLU*)^46^, and synaptic dysfunction (*PTK2B*)^47^. These findings suggest that LOAD-S1 may involve neuroinflammation and impaired clearance of toxic amyloid-beta aggregates, aligning with the cell-type enrichment patterns observed in this study. In contrast, AD genes associated with LOAD-S2 are more strongly linked to NFT pathology (i.e., *BIN1*^48^,*MAPT*^49,50^) ), amyloid plaque formation (i.e., *ADAM10*, *APP*)^50,51^, and lipid metabolism dysfunction (*ABCA7*)^52^.

The epigenomic-based LOAD subtypes identified in this study show notable correspondence with previously predicted molecular subtypes of AD. Tijms et al. categorized AD into five molecular subtypes based on CSF proteomic analysis: hyperplasticity (S1), innate immune activation (S2), RNA dysregulation (S3), choroid plexus dysfunction (S4), and blood– brain barrier dysfunction (S5)^6^. Among these, the S1 subtype closely aligns with LOAD-S2, as both exhibit a high proportion of neuronal, astrocyte, and oligodendrocyte-specific proteins, reflecting the neuronal and oligodendrocyte enrichment observed in our DNAm cell-type analysis. Additionally, we found that two out of the three categories identified in the transcriptomic-based subtypes of AD^7^ assigned to immune-related and synaptic pathway share similar characteristics with the methylomic LOAD subtypes identified in this study.

The presence of unassigned samples, which did not consistently align with LOAD-S1 or LOAD-S2 across cohorts, raises questions about their biological significance and whether they represent distinct, yet uncharacterized, subtypes, intermediate states, or result from other factors. The absence of clear cell-type-specific methylation patterns in these samples complicates their classification, limiting the study’s ability to fully capture the methylomic heterogeneity of AD. Several factors could explain the unassigned samples. They may represent mixed or intermediate phenotypes, exhibiting overlapping features of both LOAD-S1 and LOAD-S2 without strongly matching either subtype. Another possibility is the presence of co-pathologies, such as other tauopathies, alpha-synucleinopathy, or TDP-43 proteinopathies, which could influence methylation patterns. While efforts were made to screen for these co-pathologies, data availability was not uniform across all samples, potentially introducing variability that obscured subtype-specific DNAm signals. Alternatively, these samples might reflect a cohort-specific subtype, driven by unique genetic or environmental factors that are less prevalent or absent in the replication cohorts. Further investigations integrating multi-omic approaches, larger datasets, and additional neuropathological assessments will be necessary to determine whether these unassigned cases represent a distinct LOAD subtype or reflect methodological limitations. the lack of correlation between LOAD subtypes and clinicopathological markers, raising questions about how these epigenetic subtypes relate to traditional AD classifications. However, prior research has demonstrated that neuroinflammation in AD is associated with disruptions in brain network connectivity, a key driver of disease progression that occurs independently of amyloid and NFT pathology or cortical atrophy^53^. Moreover, the lack of strong correlation between transcriptomic-based AD subtypes and clinical or pathological markers in postmortem brain tissue^7^.

A key strength of this study is its large, cross-cohort design, analyzing postmortem brain samples from three independent biobanks (UKBBN, PITT-ADRC, and ROSMAP). This approach enhances the robustness of findings, ensuring validation across diverse populations while minimizing cohort-specific biases. Additionally, the study integrates genetic, transcriptomic, and cell-type-specific data, providing a comprehensive characterization of the epigenomic-based LOAD subtypes. This multi-omic approach strengthens the biological relevance of the findings and offers insight into subtype-specific molecular mechanisms underlying AD heterogeneity.

A major limitation of this study is that while it provides robust evidence of epigenetic heterogeneity in LOAD and its association with distinct biological pathways, it does not establish causality. The relationship between epigenetic heterogeneity and LOAD is likely bidirectional, with DNAm changes both contributing to and resulting from the disease process. Additionally, the lack of environmental risk factor data limits the ability to determine whether observed epigenetic changes stem from environmental exposures, lifestyle factors, or disease-related processes. The absence of detailed environmental data further restricts the ability to assess the role of non-genetic factors in driving the methylomic heterogeneity observed across LOAD subtypes. Future studies incorporating longitudinal epigenomic data and environmental risk assessments will be necessary to clarify these relationships.

The findings of this study align with previous large-scale EWASs, which have shown that DNAm changes associated with AD predominantly occur in non-neuronal cells, particularly microglia^16,33,54^. In this study, we expanded on these insights by identifying cell-type-specific DNAm signatures for the newly defined LOAD subtypes and establishing a clear link between these subtypes and distinct patterns of inflammatory microglial activity. This study highlights the importance of subtype-specific analyses in uncovering heterogeneity within complex diseases like AD and provides a foundation for future research aimed at elucidating causal pathways and identifying potential therapeutic targets. While challenges remain regarding causality and environmental influences, these findings lay critical groundwork for refining precision medicine approaches in AD research and treatment.

## Data availability

The PITT-ADRC datasets used in this study are available on Synapse (https://www.synapse.org/) under Synapse ID: syn23538600. Access requires creating a Synapse user account and submitting a data access request. The UKBBN dataset is accessible via GEO under accession number GSE284764. ROSMAP datasets are also deposited on Synapse (Synapse IDs: syn7357283, syn23650893, syn3157325). Microglial single-cell RNA sequencing data and markers of microglial states were obtained from https://compbio.mit.edu/microglia_states/. The ROSMAP mQTL dataset is accessible at https://mostafavilab.stat.ubc.ca/xqtl/. All codes used for DNA methylation and bulk transcriptomic analyses, clustering, replication, and cross-cohort validations are available at https://github.com/Dementia-Systems-Biology/LOAD_subtyping. For microglial single-cell DEG analyses, codes from https://github.com/mathyslab7/ROSMAP_snRNAseq_PFC were used.

