## Supplementary figures for "Epigenomic subtypes of late-onset Alzheimer’s disease reveal distinct microglial signatures"

**K-means clustering**

**Hierarchical clustering**

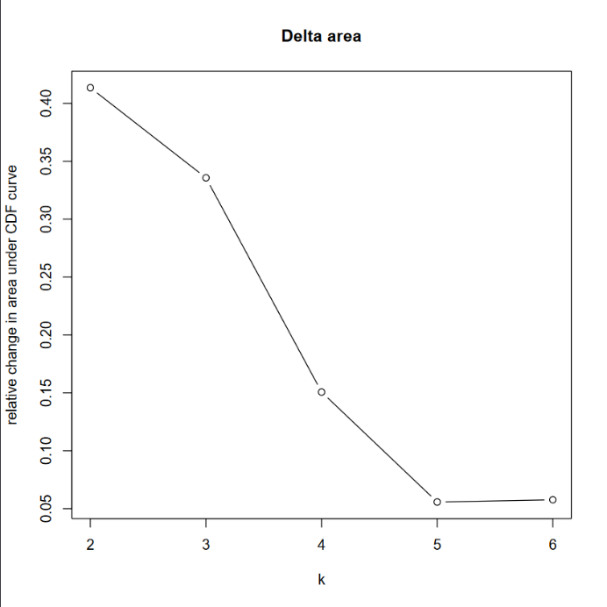

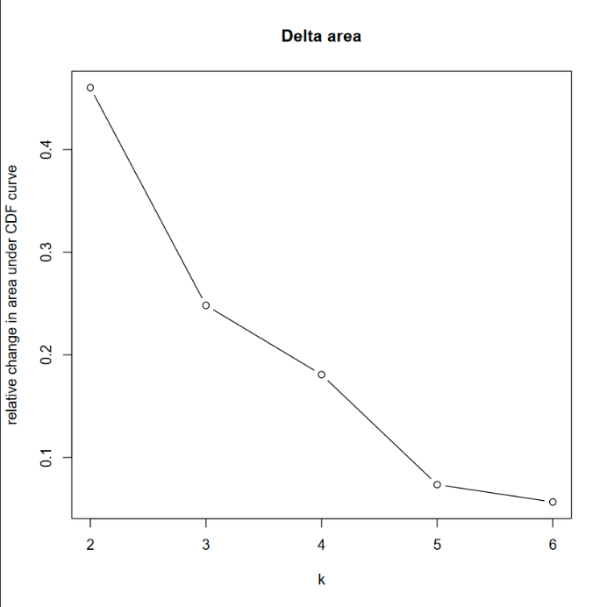

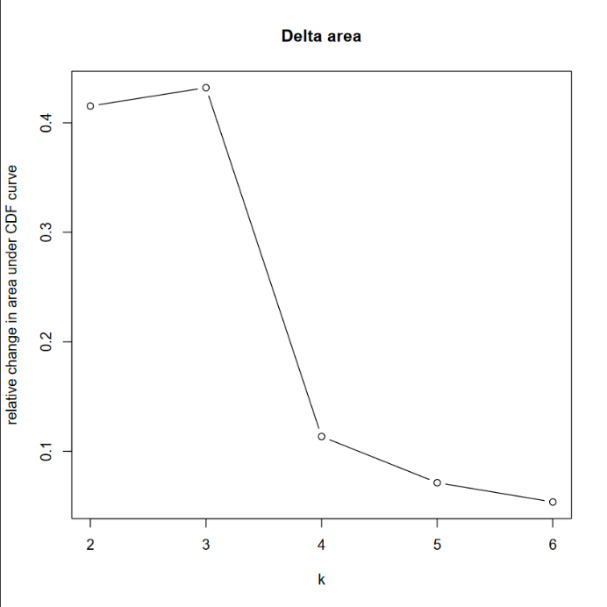

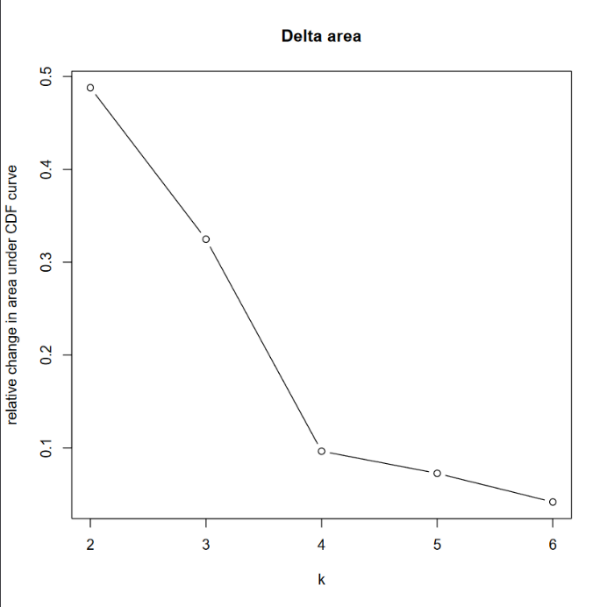

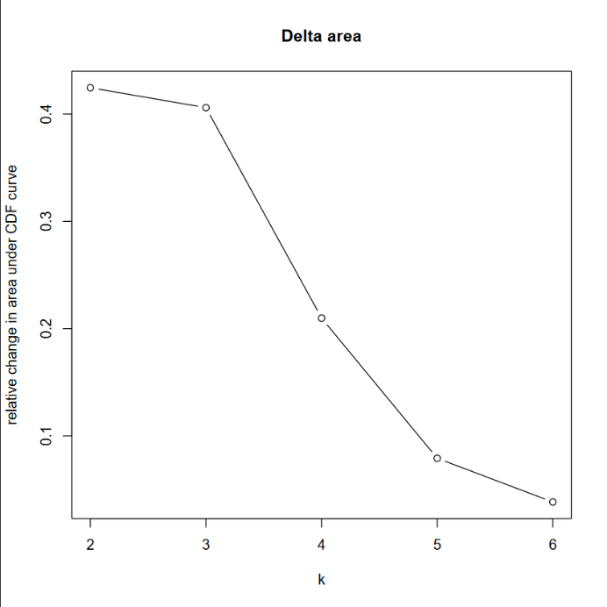

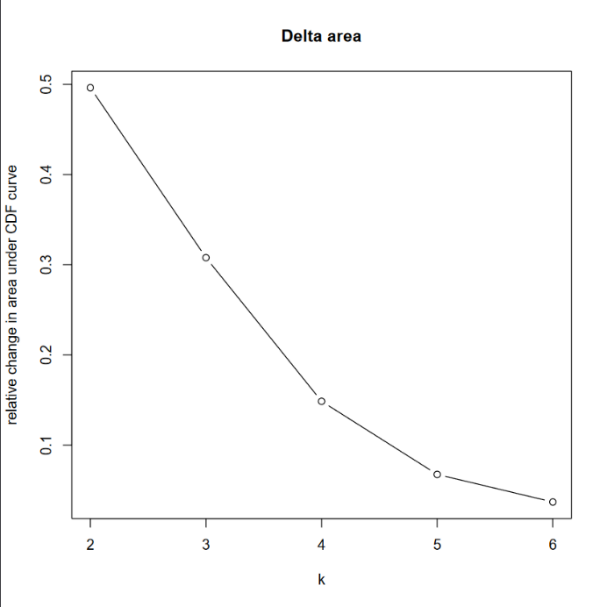

**ROSMAP**

**PITT-ADRC**

**UKBBN**

**Supplementary Figure 1. Optimal cluster determination using the Elbow method.**

Plots illustrating the application of the Elbow method to select the optimal number of clusters using Hierarchial (Left) and K-means (Right) for each cohort.

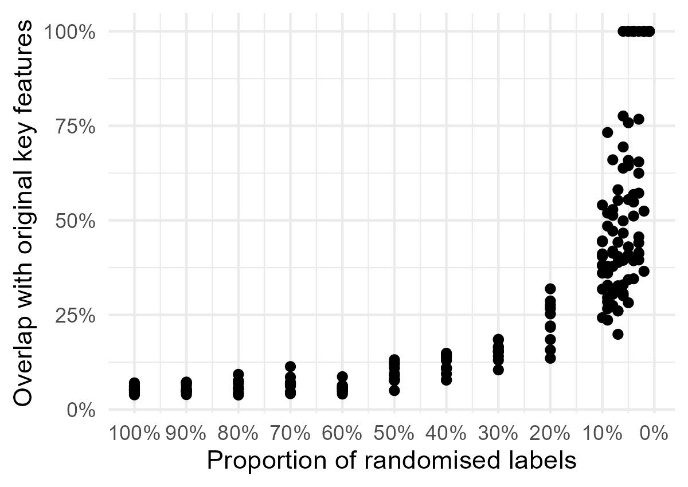

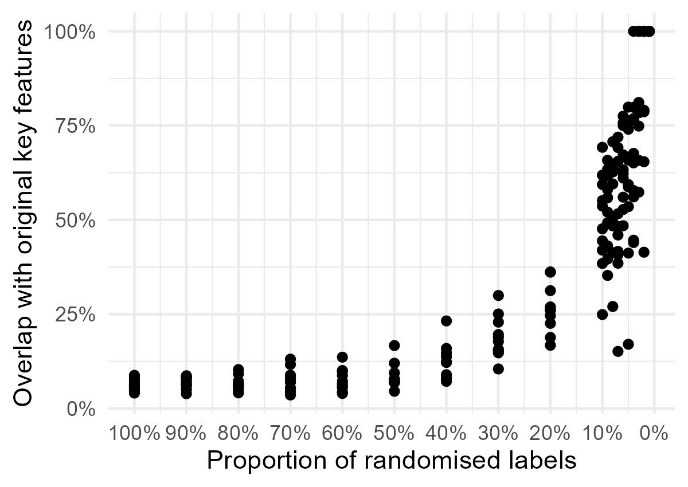

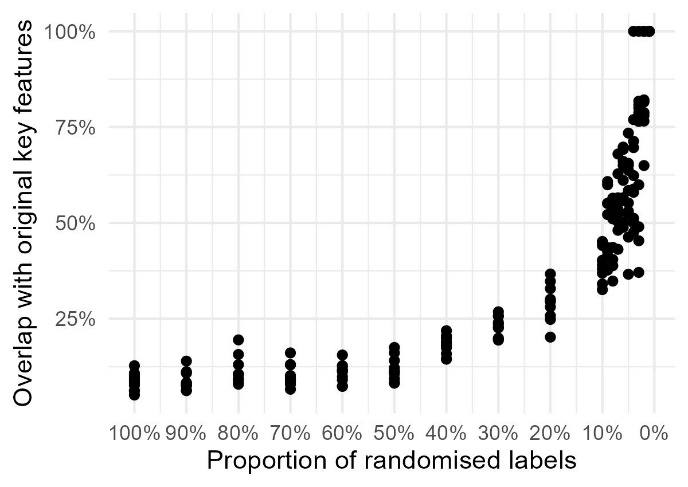

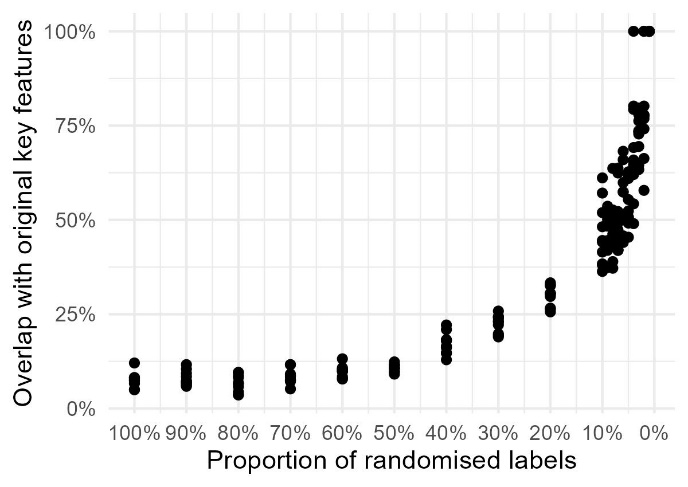

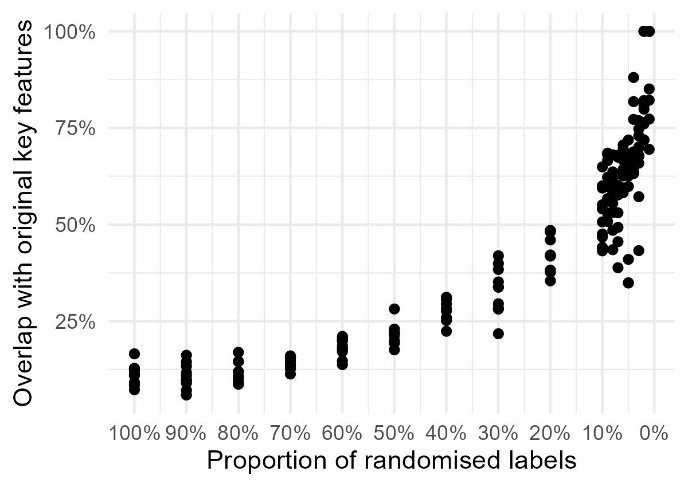

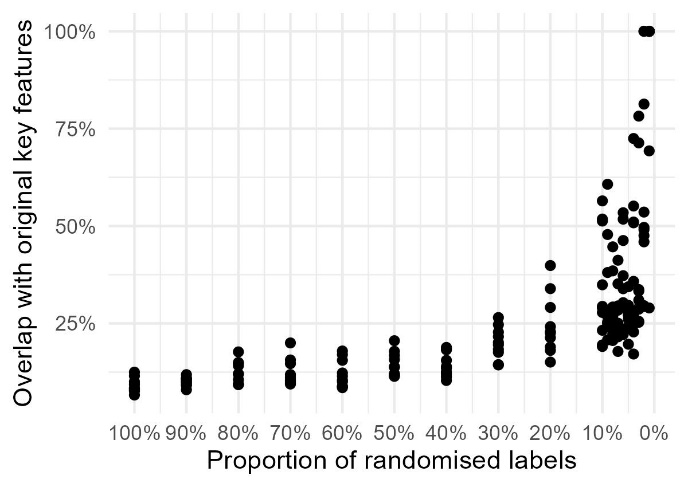

**Hierarchical clustering**

**K-means clustering**

**ROSMAP**

**UKBBN**

**PITT-ADRC**

**Supplementary Figure 2. Validation of cluster specificity through label randomization.** Clusters were validated by randomizing cluster labels across varying proportions of the sample populations. A minimal overlap between discriminative features of the original clusters and those identified using randomized labels were observed, particularly when randomization was applied to the full dataset.

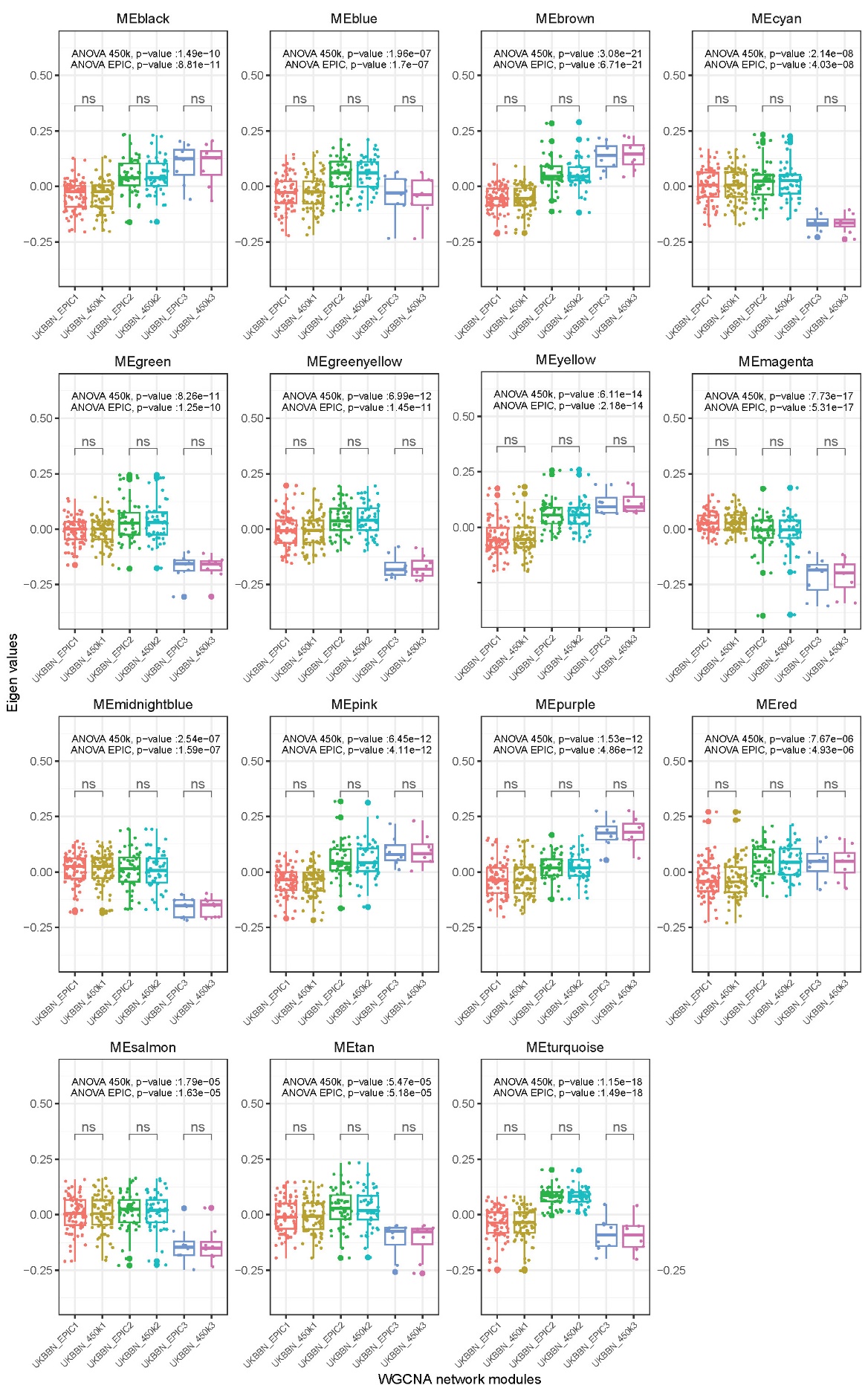

**Supplementary Figure 3. Assessment of DNAm array compatibility in the UKBBN using hierarchical clustering.** The significant relationships between module eigenvalues and the three identified clusters, as determined by ANOVA using EPIC arrays, and the subset of probes available on 450K arrays.

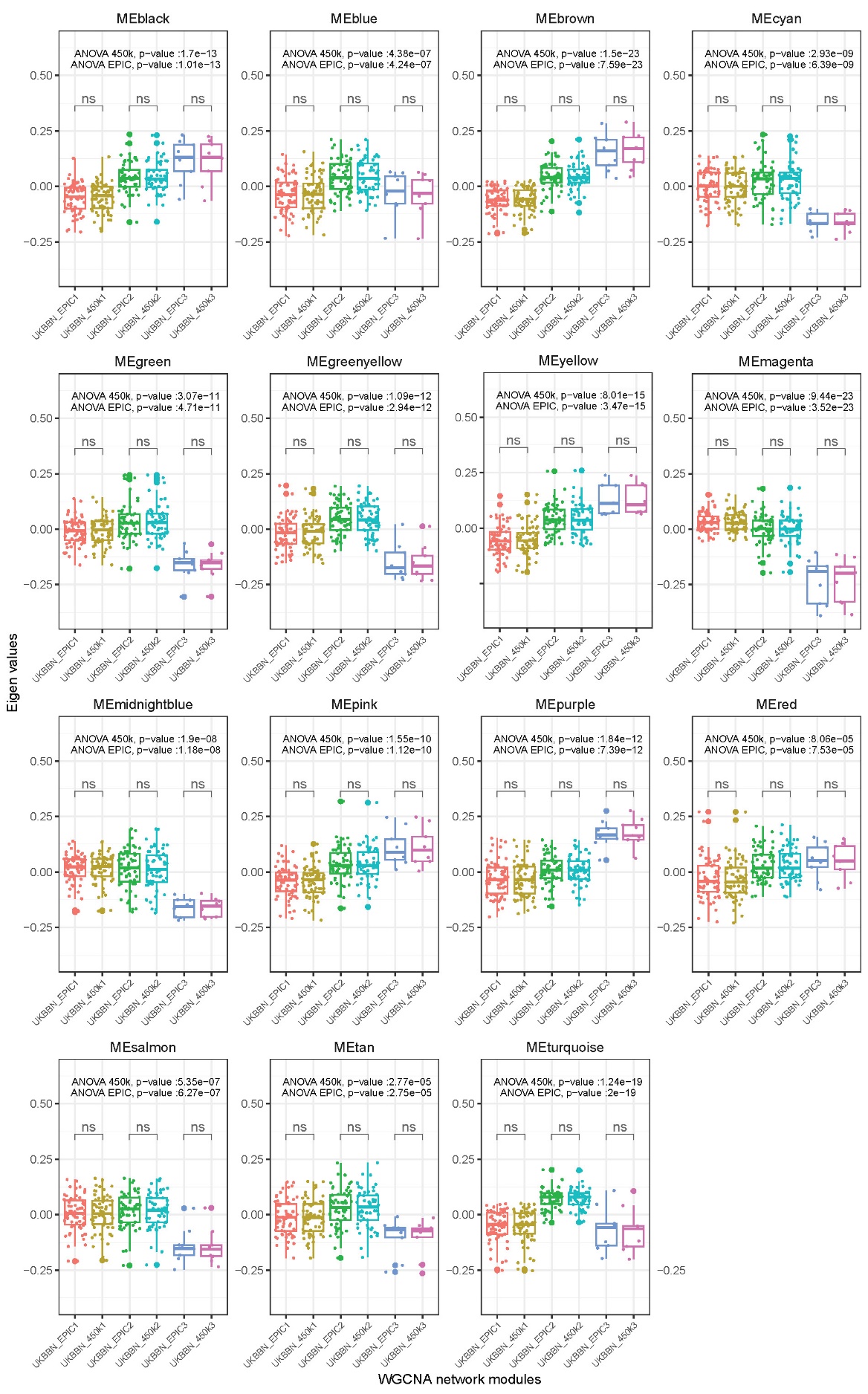

**Supplementary Figure 4. Assessment of DNAm array compatibility in the UKBBN using K-mean clustering.** The significant relationships between module eigenvalues and the three identified clusters, as determined by ANOVA using EPIC arrays, and the subset of probes available on 450K arrays.

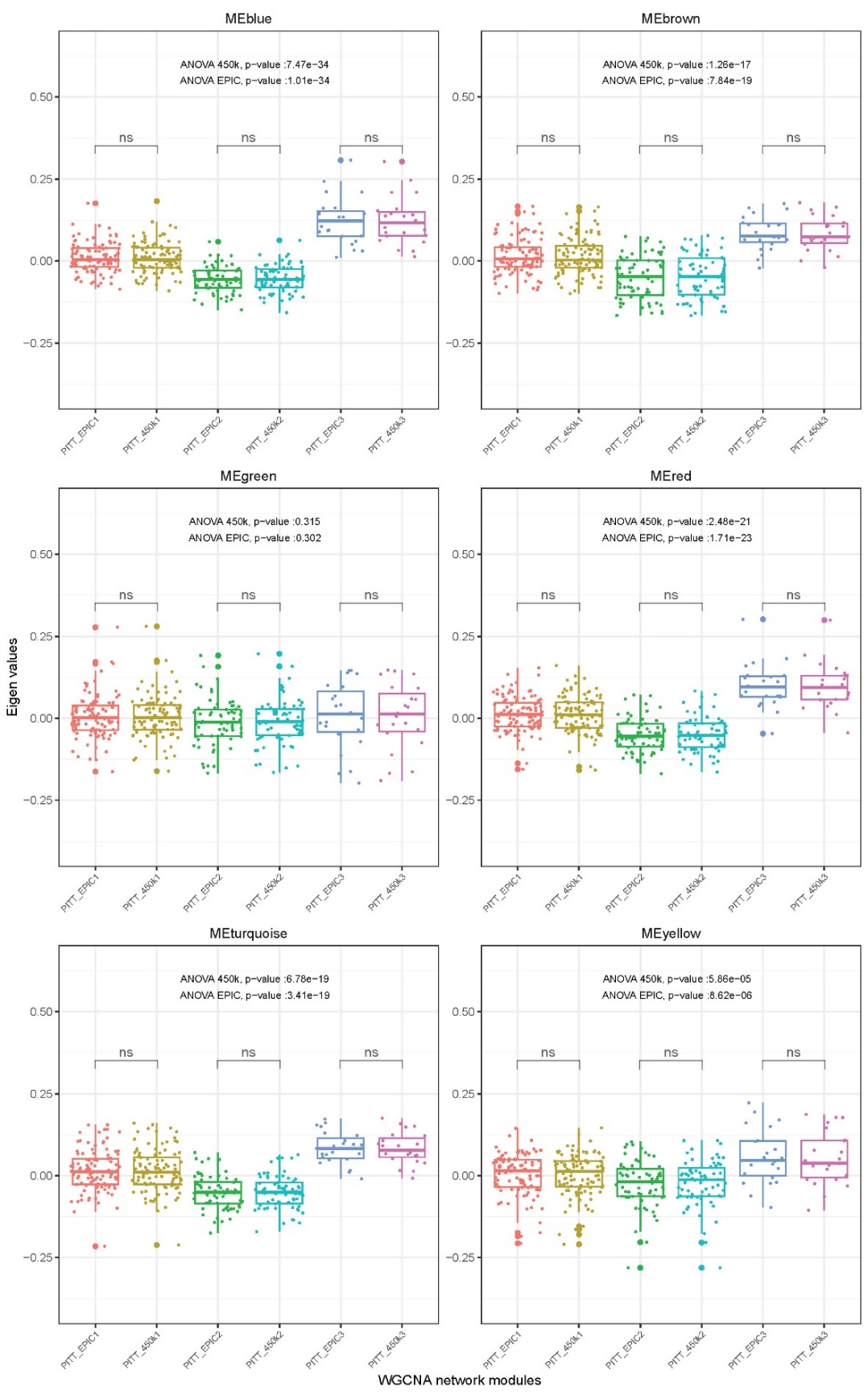

**Supplementary Figure 5. Assessment of DNAm array compatibility in the PITT-ADRC using hierarchical clustering.** The significant relationships between module eigenvalues and the three identified clusters, as determined by ANOVA using EPIC arrays, and the subset of probes available on 450K arrays.

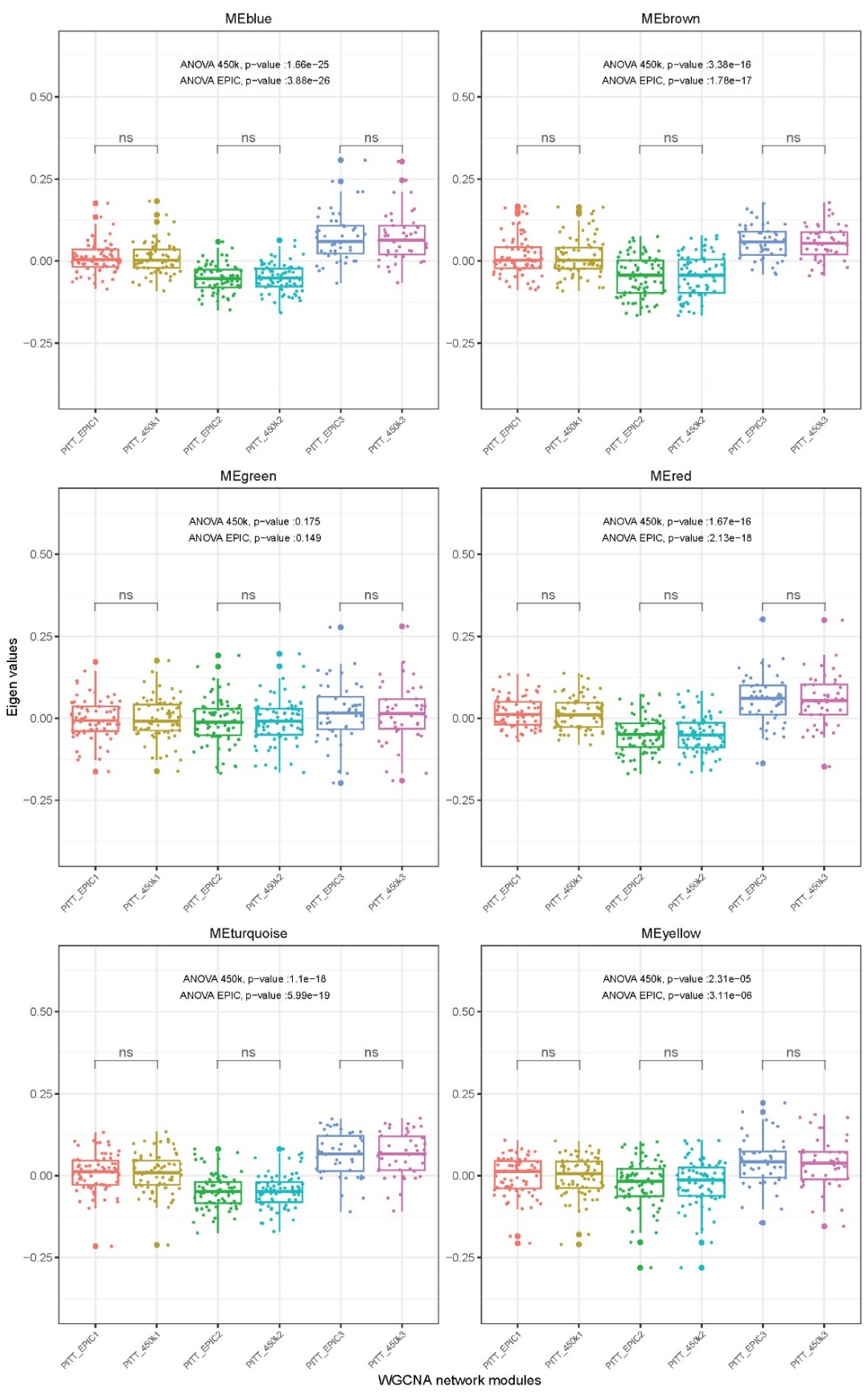

**Supplementary Figure 6. Assessment of DNAm array compatibility in the PITT-ADRC using K-means clustering.** The significant relationships between module eigenvalues and the three identified clusters, as determined by ANOVA using EPIC arrays, and the subset of probes available on 450K arrays.
